# Distinct Roles of Tumor-Associated Mutations in Collective Cell Migration

**DOI:** 10.1101/2020.06.04.135178

**Authors:** Rachel M. Lee, Michele I. Vitolo, Wolfgang Losert, Stuart S. Martin

## Abstract

Recent evidence suggests that groups of cells are more likely to form clinically dangerous metastatic tumors, emphasizing the importance of understanding mechanisms underlying collective behavior. The emergent collective behavior of migrating cell sheets *in vitro* has been shown to be disrupted in tumorigenic cells but the connection between this behavior and *in vivo* tumorigenicity is unclear. Here we use particle image velocimetry to measure a multi-dimensional collective migration phenotype for genetically defined cell lines that range in their *in vivo* behavior from non-tumorigenic to aggressively metastatic. By using cells with controlled mutations, we show that PTEN deletion enhances collective migration, while Ras activation suppresses it, even when combined with PTEN deletion. These opposing effects on collective migration phenotype of two mutations that are frequently found in patient tumors could be exploited in clinical assessments of metastatic potential or in the development of novel treatments for metastatic disease.

## INTRODUCTION

Collective migration is functionally important for metastatic dissemination, as highlighted by murine studies of multicolor primary tumors that lead to multicolor metastases through collective dissemination^1,2^. Individual phases of the metastatic cascade have also been shown to involve collective behavior. Collective invasion away from the primary tumor has been in seen in a variety of models, including intravital imaging of murine breast tumors^3^ and studies of *ex vivo* tumor spheres from colorectal cancer patients^4^. This collective dissemination from the primary tumor can lead to circulating tumor cell (CTC) clusters, which have up to 50 times higher metastatic potential than individual CTCs^5^. Additional evidence suggests that collective behavior plays a role during extravasation as well. Clusters of cells have been found to break through blood vessels intact^6^ and CTC clusters have been shown to pass through capillary-sized vessels^7^. These studies and others have identified a wide variety of collective dissemination behaviors, including multicellular streaming, epithelial collective migration, mesenchymal collective migration, and expansive growth^8^, that contribute to metastatic dissemination.

The variety of observed metastatic dissemination behaviors may be reflective of plasticity in collective cancer cell migration. This plasticity is often observed through an epithelial–mesenchymal transition (EMT) framework, in which epithelial cells exhibit decreased cell–cell adhesions, lost apical–basal polarity, and modulated cytoskeletal structures^9^. Epithelial–mesenchymal plasticity (EMP), in which cells adopt a mixture of epithelial and mesenchymal features and are potentially able to interconvert between phenotypic states^9^, has been observed in many cancer metastasis studies (as reviewed by Williams, et al.^10^). Breast cancer cells that co-express both epithelial and mesenchymal markers are more tumorigenic than cells which exhibit purely epithelial or mesenchymal phenotypes^11^. Moreover, cells exhibiting EMP are a major source of metastasis formation^12^. Increased levels of EMP were also found in CTC clusters isolated from the blood samples of triple negative breast cancer (TNBC) patients^13^. Decreases in cell–cell adhesion strength during EMP may give cells the flexibility to respond to microenvironmental cues and to transition between collective and individual migration as needed^14^. Plasticity in collective behavior is also seen through the framework of an unjamming transition, which can allow epithelial monolayers to flow collectively without necessitating the activation of EMT-transcription factors or expression of mesenchymal markers^15^. Signatures of unjamming have been observed in both non-tumorigenic MCF10A cells and in MCF10A-based models of breast cancer^16,17^. Unjamming also allows mesenchymal tumor cells to adapt to their surroundings and migrate collectively^18^.

Despite the clear importance of collective migration during metastatic dissemination, the connections between oncogenic mutations, *in vivo* tumor outcomes, and collective migration behavior are unclear. Here we use a genetically defined breast cancer model system to connect specific oncogenic mutations with a quantitative collective migration phenotype. In particular, we focus on phosphatase and tensin homologue (PTEN) loss, which is found in 24% of breast cancers^19^. The phosphatidylinositol 3 kinase (PI3K) pathway, which includes PTEN, has alterations of some type in over 70% of breast cancer patients^19^. Additionally, PTEN loss has been associated with poor outcomes in breast cancer patients^20^. Our breast cancer model system also uses overexpression of KRas(G12V), a frequent cancer driver^21,22^, alone and in combination with PTEN loss to study the response to PTEN loss in both a non-tumorigenic background and in an activated oncogenic background. The cell lines in our model system have previously characterized *in vivo* tumorigenicity^23^ and metastatic potential^24^, and have also been shown to exhibit distinct migration behavior when individual cells were allowed to move within microfluidic channels^24^. Here we develop and apply a multidimensional quantitative collective migration phenotype^25–27^ to this genetically defined breast cancer model system to measure the impact of known oncogenic mutations on collective behavior. We find that PTEN loss and activated KRas overexpression have opposing effects on migration and that the decreased collective behavior of activated KRas overexpression dominates the double mutation phenotype. Our quantitative phenotypes allow us to further investigate signatures of collective migration that are associated with the known *in vivo* tumor outcomes of our model system to determine metrics of migration that could provide insight into the tumorigenicity of a sample without requiring the selective pressures associated with long term growth in culture.

## RESULTS

We first compare the collective migration of the non-malignant MCF10A cell mammary epithelial line to the behavior of the aggressively metastatic MDA-MB-231 breast cancer cell line. Cells were plated in circular cell sheets approximately 3.6 mm in diameter (0.4 mm standard deviation). Cell sheets were imaged in regions of interest (ROIs) near the top edge and bottom edge. Example images of an MCF10A cell sheet (**Fig. 1a**) and an MDA-MB-231 cell sheet (**Fig. 1b**) are shown for the start of imaging (*t* = 0 h) and end of imaging (*t* = 12 h). To investigate features of the dynamic migration behavior of the cell sheet, images were taken every three minutes during the 12 h of time-lapse imaging, as shown in **Supplementary Movie 1.**

**Fig. 1.**
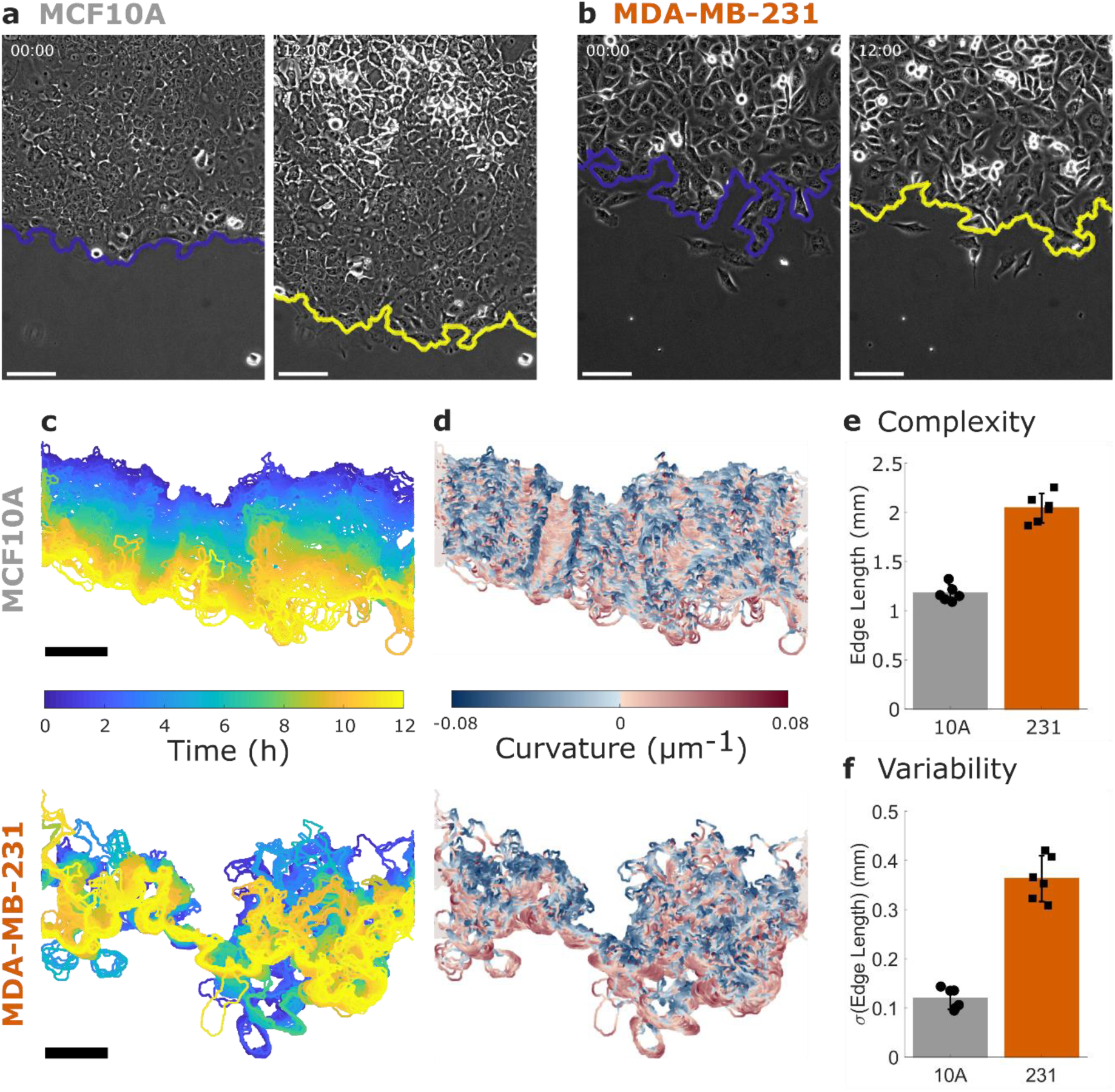
Collective Migration Changes During Cancer Progression. **a** Non-tumorigenic MCF10A cells and **b** metastatic MDA-MB-231 cells migrate in a collective migration assay over 12 hours. The leading edge of the cell sheet is indicated by a blue (initial) or yellow (12 h) line and all scale bars are 100 µm. See also **Supplementary Movie 1. c** The dynamics of the leading edge are collective in the MCF10A cells (top) but disordered in the MDA-MB-231 cells (bottom). **d** Coloring the leading edge by curvature shows persistence of local features in the MCF10A cells (top) and disorder in the MDA-MB-231 cells (bottom). **e** Edge length is used to quantify the complexity of the leading edge. **f** The variability in edge length over time is used to quantify the dynamics of the leading edge. N = 6 independent experiments. Error bars indicate 95% confidence interval.

An illustration of the cell sheet dynamics is shown in **Fig. 1c**, where the leading edge of the sheet is colored by imaging time for the example MCF10A (top) and MDA-MB-231 (bottom) cell sheets shown in **Fig. 1a-b**. The smooth transition from blue (0 h) to yellow (12 h) indicates that the leading edge of the MCF10A sheet progresses steadily forward, while the MDA-MB-231 leading edge shows disordered progression over time. Coloring each edge by time (**Fig. 1c**) indicates the displacement of the edge over time, while coloring the leading edge by the local curvature of the cell sheet (**Fig. 1d**) emphasizes the persistence of local features. In the MCF10A sheet, persistent regions of blue and red (i.e. high curvature regions) are seen to progress forward over time, while the MDA-MB-231 sheet does not show consistent features. As the sheets are all imaged in a ROI of the same width, the length of the leading edge reflects the complexity of its bends and turns across the image. Comparing the average edge length between the MCF10A and MDA-MB-231 cells (**Fig. 1e**), we find that the MDA-MB-231 edge is on average 0.9 mm longer than that of the MCF10A. Compared to the minimum possible edge length of approximately 0.6 mm (the width of the imaging ROI), this increase of 0.9 mm indicates a substantial increase in complexity in the MDA-MB-231 leading edge. We also investigate the variability of the edge length over time (**Fig. 1f**) and find that the MDA-MB-231 cells have approximately three times the variability of the MCF10A cells.

To further analyze the dynamics of migration in the cell sheets, we use Particle Image Velocimetry (PIV) to measure flows within the cell sheet. Example PIV flow fields for the MCF10A and MDA-MB-231 cells are shown in **Fig. 2a**. We note that these flow fields capture the motion of the entire cell sheet, including subcellular motion. Overall flow speeds calculated from PIV do not show differences between the MCF10A and MDA-MB-231 cells (**Fig. 2b**), despite the differences in leading edge displacement illustrated in **Fig. 1** and quantified in **Fig. 2c**.

**Fig. 2.**
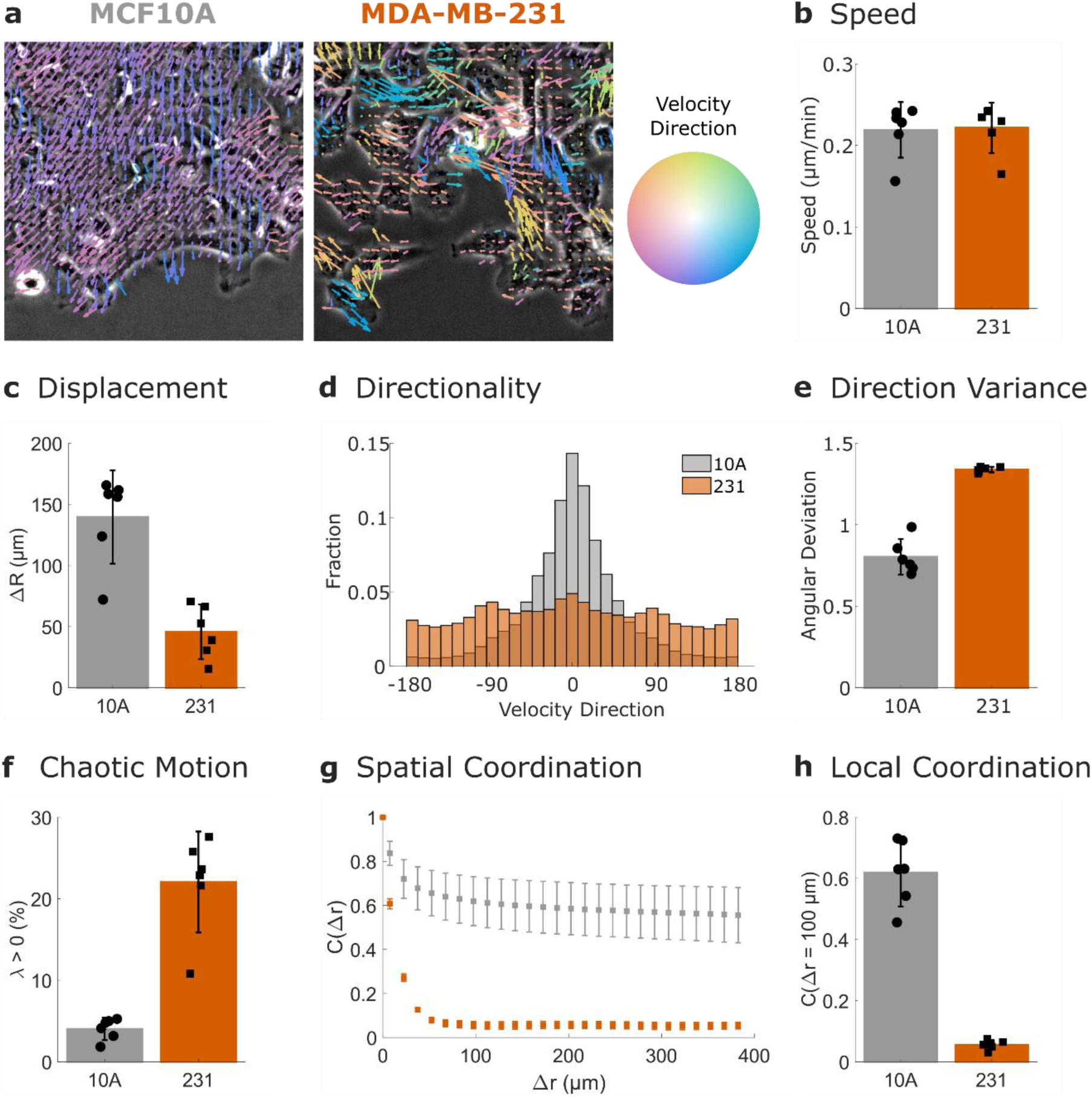
Collective Behavior Quantitatively Decreases in Metastatic Cells. **a** PIV flow vectors are colored by the direction of motion and overlaid on images of MCF10A cells (left) and MDA-MB-231 cells (right). **b** Mean speed of the PIV flow field. **c** Displacement of the leading edge. **d** Distribution of velocity vectors. **e** Variability in velocity direction quantified by angular deviation. **f** Chaotic motion in the cell sheet quantified by the percentage of positive finite-time Lyapunov exponents (λ). **g** Spatial coordination across multiple length scales quantified by the spatial autocorrelation of the radial velocity. **h** Local coordination is defined as the spatial autocorrelation value at 100 µm (a few cell lengths). N = 6 independent experiments. Error bars indicate 95% confidence interval.

This difference is reconciled by measurements of the directionality within the cell sheet. As shown by the color of the flow field vectors in **Fig. 2a**, the MCF10A cells mainly flow into the empty space at the sheet edge, while the MDA-MB-231 cells migrate in multiple directions. Cumulative distributions of velocity direction across six biological replicates (**Fig. 2d**) emphasize the strong differences in directionality between the two cell lines. This is further quantified by the angular deviation of these distributions (**Fig. 2e**); the higher values in the MDA-MB-231 cells indicate less directional migration.

Additional metrics of collective migration can provide insight into which features of migration are most affected by perturbations^25,27^, thus we also measured the behavior of the MCF10A and MDA-MB-231 cell lines over long length and time scales. The directionality in **Fig. 2d-e** is measured on the scale of the imaging frame rate (minutes). Finite time Lyapunov exponents (λ), however, are a measure of the divergence of nearby trajectories in the flow over longer time scales (in this case, 2 h). Positive values of λ are associated with chaotic flows and thus increased percentages of positive λ values (**Fig. 2f**) indicate decreased coordination in the MDA-MB-231 cells over long timescales.

To measure the coordination of the cell sheet over long distances, we compute the spatial autocorrelation of the PIV flow field’s radial velocity (**Fig. 2g**). The MCF10A cells show similarity with neighbors hundreds of µm away, while the MDA-MB-231 cells show decreased correlations after approximately 50 µm. To quantify the coordination of cells with their local neighbors, we compare the autocorrelation values at 100 µm in **Fig. 2h**, which shows higher spatial coordination in the MCF10A cell line.

Our finding that the MDA-MB-231 cells exhibit a less collective migration phenotype than the MCF10A cells is in agreement with previous studies which have found disordered collective behavior in cancer cell types^16,28^. Next we investigated the collective behavior of a genetically defined cancer model system which has been previously characterized *in vivo* for both primary tumorigenesis and metastatic efficiency^23,24^. This comparison allows us to connect our multidimensional collective migration phenotype to mutations with known tumor outcomes. This model system is based on the MCF10A cell line, which does not form tumors in murine models. Deletion of PTEN in MCF10A cells allows the cells to remain dormant *in vivo*, while activation of KRas in the MCF10A leads to dormancy or, rarely, to the formation of tumors^23^. However, when these two mutations are combined, KRas/PTEN^-/-^ cells are able to form aggressive primary tumors^23^ and metastasize in the lungs^24^. Images of this model system in the collective migration assay are shown in **Fig. 3a** and in **Supplementary Movie 2**. As seen in **Fig. 3a** and quantified in **Supplementary Fig. 1**, the cell lines with overexpressed active KRas show increased complexity in their leading edge.

**Fig. 3.**
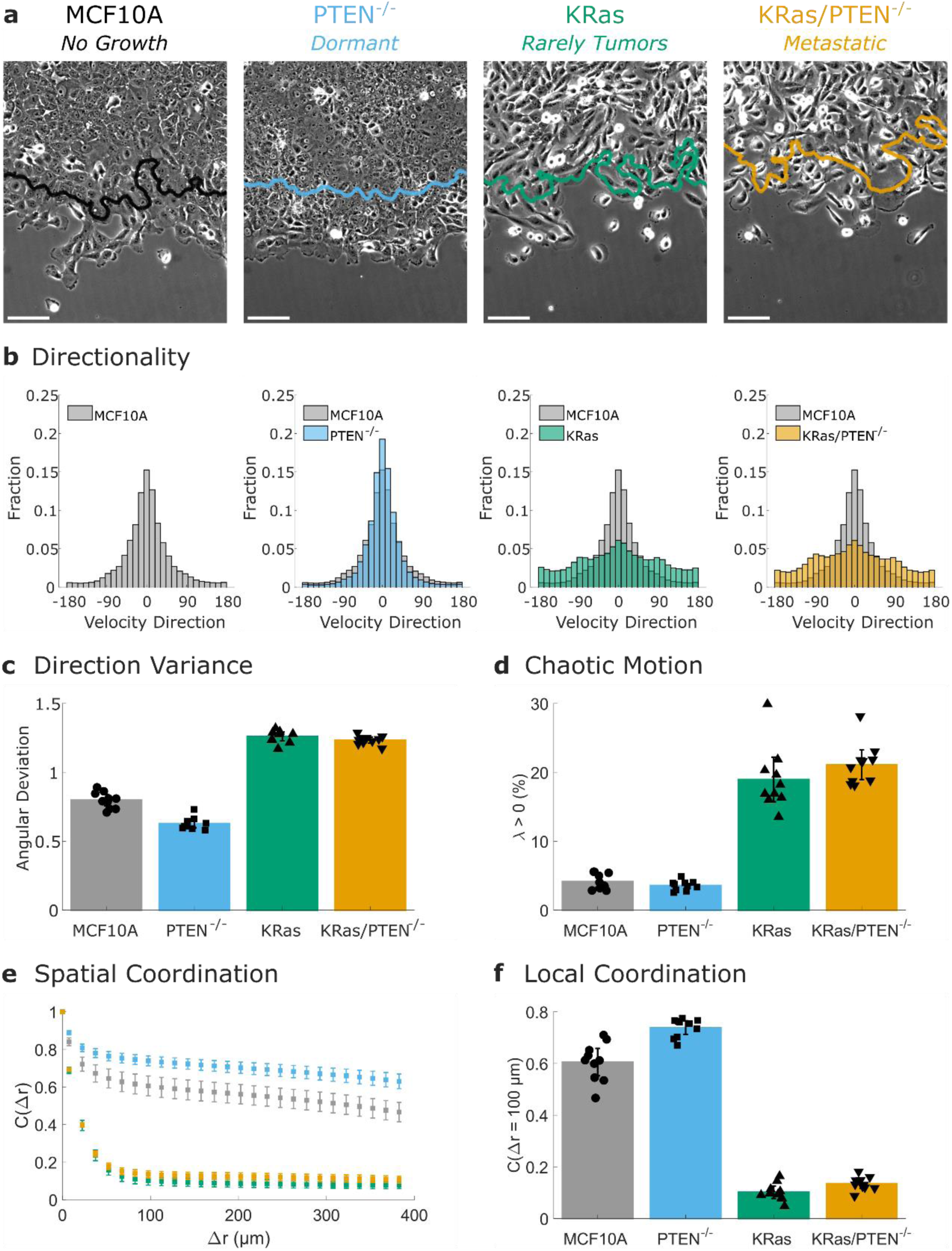
Activated KRas and PTEN^-/-^ have opposing effects on collective migration. **a** Images from collective migration assays on cells from a genetically defined cancer model system after 12 h of migration. The position of the initial (t = 0 h) leading edge is indicated by a line on each image and all scale bars are 100 µm. See also **Supplementary Movie 2. b** Distributions of velocity direction for each cell line are compared to the non-tumorigenic MCF10A control. **c** Variability in velocity direction quantified by angular deviation. **d** Chaotic motion in the cell sheet quantified by the percentage of positive finite-time Lyapunov exponents (λ). **e** Spatial coordination across multiple length scales quantified by the spatial autocorrelation of the radial velocity. **f** Local coordination is defined as the spatial autocorrelation value at 100 µm (a few cell lengths). N = 10 independent experiments. Error bars indicate 95% confidence interval.

PIV analysis of these cells lines indicates that the PTEN^-/-^ cells are slightly more directional than the MCF10A control, while both the KRas and KRas/PTEN^-/-^ cells are less directional, as shown by the cumulative direction distributions (N = 10 biological replicates) in **Fig. 3b**. The KRas and KRas/PTEN^-/-^ cells both show increased angular deviation values (**Fig. 3c**) compared to the MCF10A, in a similar manner as the MDA-MB-231 cells (**Fig. 2e**). This increase in angular deviation is reflected in the decreased progression of the leading edge as shown by the edge overlays in **Fig. 3a** (quantified in **Supplementary Fig. 2b**). The KRas and KRas/PTEN^-/-^ cells also show increased chaotic motion compared to the MCF10A control (**Fig. 3d**) as well as decreased spatial coordination over long distances (**Fig. 3e-f**). The PTEN^-/-^ cells show the opposite trend of increased coordination over long distances and thus trend towards increased collective behavior in multiple metrics. The double mutant KRas/PTEN^-/-^ cells are dominated by the effect of the KRas mutation, which shows decreased collective behavior on time scales of minutes and hours, as well as across long distances.

To compare multiple metrics with varied scales and units of measurement, we convert the results shown in **Fig. 1-3** into paired t-statistics (see **Supplementary Fig. 3**). The multidimensional collective migration phenotype can then be summarized as shown in **Fig. 4a**. Migration metrics, shown on the y-axis, are colored by whether the metric would be expected to decrease (blue) or increase (red) as collective behavior increases. The four cell lines, shown on the x-axis, are compared to the MCF10A cell line as a control. **Fig. 4a** shows that while the PTEN^-/-^ cells tend to increase in collective behavior, the KRas, KRas/PTEN^-/-^, and MDA-MB-231 cells show decreased collective behavior. In **Fig. 4b**, the KRas/PTEN^-/-^ cells are compared to the individually mutated cell lines; the higher similarity with the KRas individual mutation cell line emphasizes the dominant nature of KRas on the collective migration phenotype.

**Fig. 4.**
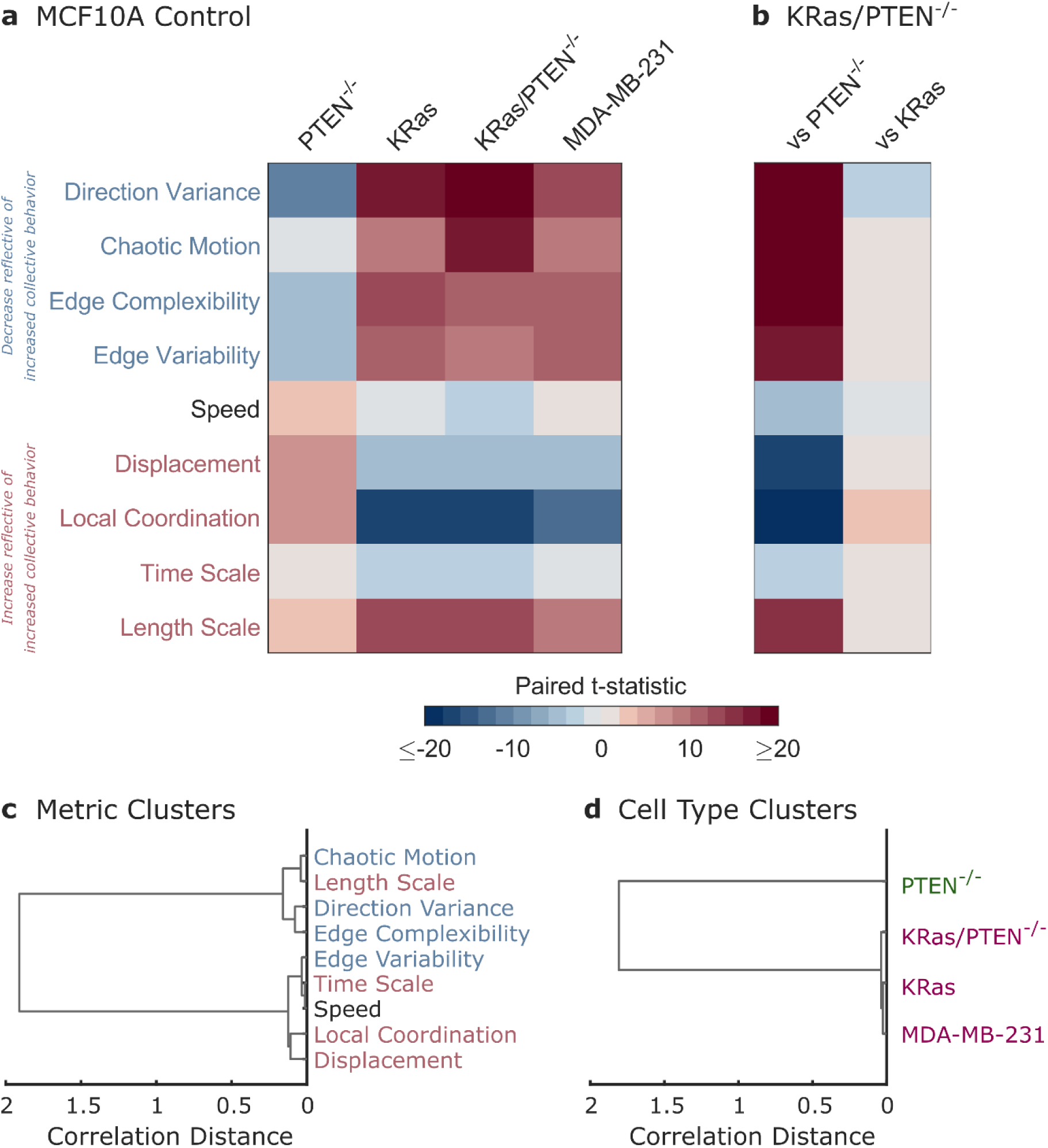
Activated KRas dominates the collective migration phenotype. **a** Summary of the multidimensional phenotype for four cell lines as compared to the paired MCF10A control. The strength of changes in migration metrics are indicated by color representing the paired t-statistic (see **Supplementary Fig. 3**). **b** Summary of migration phenotypes comparing the double mutant KRas/PTEN^-/-^ cells to the single mutant cell lines. **c** Dendrogram showing clustering of migration metrics using correlation distance. **d** Dendrogram showing clustering of cell lines using correlation distance.

Clustering of the migration metrics based on correlations within the cell lines (**Fig. 4c**), reveals that the metrics largely separate as would be expected into groups which either increase (red) or decrease (blue) as collective behavior increases. One deviation from this trend, the characteristic length scale of motion, could reflect both changes in cell size as well as changes in spatial coordination. Clustering of the cell lines combined with fitting the data into distinct groups leads to two clusters: the highly collective PTEN^-/-^ cells and the three cell lines with disordered collective migration (**Fig. 4d**).

## DISCUSSION

Using a multidimensional, quantitative collective migration phenotype, we find that two tumor-associated mutations (PTEN loss and activated KRas overexpression) have distinct effects on behavior. We find that PTEN deletion leads to an increase in collective behavior compared to the MCF10A control cell line. In contrast, KRas(G12V) overexpression leads to a clear decrease in collective behavior, similar to the decrease in collective behavior observed in the highly metastatic MDA-MB-231 cell line. The double mutant KRas/PTEN^-/-^ cell line also shows decreased collective behavior, suggesting that KRas dominates the collective migration phenotype and that the effects of both mutations are not additive. This result is in agreement with the known KRas mutation (G13D) in the MDA-MB-231 cell line^29^, which is often used a breast cancer model system for rapid metastasis. Although Ras mutations are thought to be rare in breast cancer, KRas is one of the most frequently altered genes across tumor types^21,22^, persistent activation of Ras is seen in many breast cancer cell lines^30^, and components of the Ras signaling pathway (e.g. growth factor receptors or ERK) are mutated or highly activated in breast cancer^31,32^. This signaling pathway has also been implicated in collective dissemination. A recent study of an MCF10A-based model of unjamming in which the small GTPase RAB5A is overexpressed lead to collective invasion in a 3D acini model^17^. This result was mediated by ERK1/2 activation, supporting the idea that pathways downstream of KRas activation may play an important role in collective breast cancer dissemination.

Although KRas dominates the measured collective migration phenotype, prior work shows that this mutation alone is not sufficient for aggressive tumorigenicity^23^ or metastasis^24^ *in vivo. In vitro* studies have also found that Ras activation alone is not sufficient for breast cancer cell lines to have invasive properties^30^. The trend toward increased collective behavior observed in the PTEN^-/-^ cells raises the possibility that the opposing migratory phenotypes of PTEN^-/-^ and KRas may provide additional plasticity to the double mutant line and enable EMP in response to *in vivo* microenvironments. The cancer microenvironments encountered during collective dissemination can include changes in stiffness or texture that are not captured by our *in vitro* collective migration assay^33,34^. Prior work has shown that cells that have undergone an unjamming transition or that exhibit EMP are able to migrate collectively in response to *in vivo* microenvironmental cues^14,18^. The flexibility provided by EMP may be especially relevant during the circulation phase of metastasis, where EMP signatures such as E-cadherin expression are required for successful collective dissemination^35^. PTEN loss may thus allow cells overexpressing KRas the additional flexibility necessary to adapt their collective behavior in response to *in vivo* cues.

PTEN loss may also impact other essential pathways for successful metastasis. Migration and invasion have been recognized as a central hallmark of cancer^36^, but there are many other malignant transformations that contribute to cancer progression^37^. Prior work has shown that PTEN loss leads to growth-factor independent proliferation, as well as likely resistance to apoptosis^38^. The apoptotic insensitivity conferred by PTEN loss may play an important role in our breast cancer model system in light of previous results showing that apoptotic resistance does not directly promote tumor growth, but can increase metastasis when combined with activation of the Ras-MEK pathway^39^. The need for additional information to distinguish the metastatic potential of the KRas and KRas/PTEN^-/-^ cell lines is also in agreement with the observed individual migration of these cell lines. Individual cell migration within a microfluidic channel was found to be highly predictive of metastatic potential, but adding Ki-67 as a marker of proliferating cells allowed for the removal of false positives and helped distinguish between the KRas and KRas/PTEN^-/-^ cell lines^24^.

Markers such as Ki-67 are already used clinically for breast cancer prognosis. Increasing evidence suggests that migration may be more predictive of outcome than cell growth^24,40^. Quantitative migration phenotypes have also been used to identify novel targets of dissemination that may not have been identified by screens of growth alone^41^. This suggests that adding assessments of collective migration behavior such as the quantitative phenotype described here may allow for more accurate clinical assessments of metastatic potential. The simple collective migration assay employed here does not require cell growth and thus avoids the selective pressures associated with long-term culture or expansion of patient tumor samples *in vitro* or in mouse models. This assay also uses relatively simple phase-contrast microscopy and does not require fluorescent labeling of the tumor sample. The 12 h time course analyzed in this work allows for the measurement of metrics on multiple length and time scales that can help distinguish collectively migrating cell lines which behave similarly on shorter time scales^25^. As metastasis occurs over days and weeks *in vivo*, measurements of dynamic migration behavior over hours add depth to cancer phenotypes. However, several of the metrics in our multidimensional collective migration phenotype rely on measurements on the scale of minutes (direction variance, edge complexity, speed, local coordination, and length scale) and could thus be applied to shorter series of time lapse images to allow for faster assessment of collective migration behavior.

Our multidimensional collective migration phenotype integrates migration data on a wide range of length and time scales, and thus captures effects from a broad range of biophysical processes. Given the importance of migratory phenotypes for metastatic potential, our analysis could be incorporated into future tools to provide rapid assessments of patient tumor cells on the scale of several hours, compared to the days to weeks required for spheroid growth assays or the months required for PDX models. As expected, cancer metastasis is a complex phenotype and collective migration assessments alone are not sufficient to capture the differences in metastatic potential of the KRas and KRas/PTEN^-/-^ cells. We propose that PTEN loss could drive additional hallmarks of cancer which could be assessed in combination with collective migration phenotypes. For example, enhanced cell proliferation may be detected with clinical markers, such as Ki-67^24^, while enhanced plasticity may be directly tested through *in vitro* microenvironmental cues, such as texture^42,43^. Quantitative assessments of collective migration also allow for the investigation of the role of specific oncogenic mutations in collective dissemination as well the impact of drug treatments on collective migration^27^. This creates opportunities for future studies to identify potential targets of collective dissemination *in vivo* and to also identify the role current treatments may play in curtailing or unintentionally enhancing^44^ metastatic progression.

## METHODS

### Cell Culture

MCF10A cells (ATCC) were maintained in DMEM/F12 medium (Thermo Fisher Scientific 11-330-057) with 5% horse serum (Thermo Fisher Scientific 26050088) that was supplemented with 10 μg/ml insulin (ThermoFisher Scientific 12585014), 10 ng/ml EGF (PeproTech AF-100-15), 0.5 μg/ml hydrocortisone (Sigma-Aldrich H4001), and 100 ng/ml cholera toxin (Sigma-Aldrich C8052). MDA-MB-231 GFP-LUC cells^23,24^ were maintained in DMEM (Corning 10-017-CV) with 10% FBS (Gemini Bio-Products 100-106). MCF10A, MCF10A PTEN^-/-38^, MCF10A KRas(G12V)^23^, and MCF10A KRas(G12V)/PTEN^-/-23^ cells were infected with a LifeAct-GFP expressing lentivirus; LifeAct cells were used in all comparisons between these four cell lines reported in this work. All LifeAct cells were cultured in MCF10A media supplemented with 0.5 μg/ml puromycin (AG Scientific P-1033-sol). All cells were maintained in a humidified atmosphere at 37 °C and 5% CO_2_. All cells used in this study tested mycoplasma negative using the Lonza MycoAlert (VWR 75870-454) testing system.

### Migration Assay and Imaging

Glass bottom 12-well plates (MatTek Corporation P12G-1.5-14-F) were coated with 3.25 μg/cm^2^ collagen IV (Corning 354233). The plate was placed on ice and 500 ml of collagen IV solution (3.25 μg/cm^2^ collagen IV in 0.05 M HCl) was added to each well for 1 h before rinsing each well twice with water. Cells for the migration assay were suspended at a concentration of 1.5 × 10^6^ cells/ml and plated as a 5 µL drop in the center of each well. After approximately 45 min at 37 °C, non-adherent cells were removed by medium rinses, the well was filled with 1 ml medium, and the cells were allowed to adhere in the incubator overnight. This resulted in a roughly circular sheet of cells surrounded by empty space.

Approximately 1 h before the start of imaging, the medium in each well was replaced with 1 ml of fresh medium. Cells were imaged overnight on a PerkinElmer spinning-disk confocal microscope with a 10× phase-contrast objective (0.582 µm/pixel). Images were collected using PerkinElmer’s Volocity software (version 6.4.0) with a Hamamatsu C10600-10B ORCA-R2 camera that recorded 12-bit images. Images were recorded every 3 min for 12 h.

### Image Analysis

The leading edge of each migrating cell sheet was segmented using custom MATLAB code as previously described^25^. Sobel filtering, median filtering, and morphological operators were used to find a binary image that indicated the location of the cell sheet. Dijkstra’s algorithm, as implemented by Sebastien PARIS for MATLAB^45^ was used to find the coordinates of the leading edge. Two opposing edges of each cell sheet were imaged over time; these two edges were fit to a circle to calculate the radius of the approximately circular cell sheet over time. The two edges combined with the microscope stage positions from overnight imaging were used to create a polar coordinate system for each cell sheet to define the radial direction of motion (the direction of motion expanding the sheet) as previously described^25^.

Particle image velocimetry (PIV) analysis was based on the MatPIV toolbox by Kristian Sveen (GNU general public license)^46^. This toolbox was used to extract velocity information from the time-lapse images as previously described^25,26^. Briefly, two first-pass calculations with 64 pixel × 64 pixel windows (approximately 37 µm × 37 µm) were followed by two iterations of 32 pixel × 32 pixel windows, in all cases with a 50% overlap. Outliers were detected using a signal-to-noise filter (threshold of 1.3).

### Migration Analysis

The length of the segmented cell sheet edge was calculated as the summed distance between all boundary points. As all leading edges spanned the same image width, the mean edge length over time was used as a measure of the edge complexity. The standard deviation of the edge length over time was reported as the edge variability. The difference between the calculated cell sheet radius at *t* = 0 h and *t* = 12 h was reported as the cell sheet displacement. Curvature of the edge was calculated as the Menger curvature of points within 40 boundary points of the boundary point of interest.

Bar graphs of speed report the mean speed of PIV vectors averaged over both time and space. Direction distributions indicate the direction of PIV vectors with respect to the radial direction of motion and are shown as cumulative distributions across biological replicates. The variability in velocity direction was parametrized by the angular deviation, calculated as 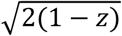 where *z* is defined in **equation (1)**. This equation bounds the values of angular deviation between zero (no variance in directionality) and 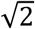 (highly varied directionality).

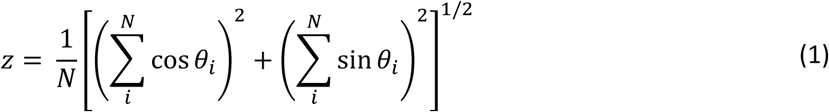

Finite-time Lyapunov exponents (λ) were calculated by computationally tracing virtual particles through the PIV flow field. As previously described, we use a deformation time of 2 h^26^ to allow the λ values to asymptotically approach Lyapunov exponents^47^. Tracer particles are initiated on the PIV grid but allowed to move off the grid as the flow field evolves over time. For each set of four virtual particles that were initially neighbors, the logarithm of the largest eigenvalue of the Cauchy–Green strain tensor is the local λ value. Positive λ values indicate an exponential sensitivity to initial conditions and are associated with chaotic flow fields, thus the percentage of positive λ values is used as a measure of the chaotic motion in the system.

Spatial autocorrelations were calculated for the radial velocity component of the PIV flow field, *v*, using **equation (2)**. The value of the autocorrelation at Δ*r* = 100 µm was used as a metric of local coordination within the cell sheet.

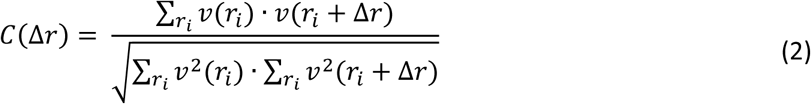

To calculate characteristic length and time scales of motion from the PIV flow fields, we used a coarse graining technique described by Zorn, et al^48^. By averaging the PIV flow field over increasing time intervals, the variance of the flow field decreases. This variance can be fit to an exponential decay as shown in **equation (3)** to calculate the characteristic time scale *t*_*c*_. The second term of this equation accounts for the finite accuracy of the microscope stage in returning to a position during multi-position imaging (*t*_*imaging*_ = 3 min).

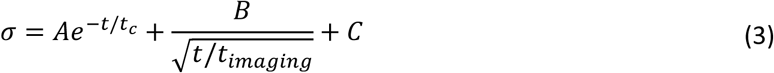

Similarly, the characteristic length scale of motion was calculated by averaging the PIV flow field over increasing spatial distances. The decreasing variance in the flow field can then be fit to the exponential decay in **equation (4)** to calculate the characteristic length scale *l*_*c*_.

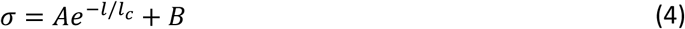

### Clustering Analysis

Migration assay imaging data was collected as paired biological replicates where multiple cell lines were imaged in the same 12-well plate over time using multi-position time-lapse imaging. To compare multiple metrics in a multidimensional collective migration phenotype, paired t-statistics were calculated for each metric. As shown in **Supplementary Fig. 3**, a paired difference was first calculated for each cell line compared to the control. The mean of this paired difference divided by the standard deviation of the paired difference is the paired t-statistic, which is used to report summaries of multidimensional migration phenotypes in **Fig. 4**.

Clustering of cell lines and migration metrics was based on the correlation distance (one minus the sample correlation between points). An agglomerative hierarchical cluster tree was calculated using MATLAB’s `linkage` function using the ‘average’ method for the ‘correlation’ distance metric between clusters. Dendrograms in **Fig. 4** were created using the leaf ordering determined by MATLAB’s `optimalleaforder` function.

The cluster tree was divided into distinct clusters as previously described^27,49^. Clustering matrices for the metrics and cell lines respectively were ordered according to the optimal leaf order. This matrix was then fit to a block-diagonal structure using greedy algorithm for splitting the data into blocks and a Bayesian information criterion (BIC) to avoid overfitting in determining cluster boundaries.

## DATA AVAILABILITY

Source data for Fig. 1e-f, Fig. 2b-h, Fig. 3b-f, Fig. 4a-b, Supplementary Fig. 1c-d, and Supplementary Fig. 2-3 are available in Supplementary Data. Additional data from the current study are available from the corresponding author on request.

## ACKNOWLEDGEMENTS

This research was supported by the METAvivor foundation and NIH grants T32-CA154274, R01-CA154624 and R01-CA124704 as well as AFOSR grant number FA9550-16-1-0052, Veterans Administration Merit Award I01-BX002746, and the American Cancer Society Research Scholar Grant RSG-18-028-01-CSM. We thank the University of Maryland Imaging Incubator for the use of their spinning disk microscope for this project. The Marlene and Stewart Greenebaum Comprehensive Cancer Center is supported by P30-CA134274 and the Maryland Cigarette Restitution Fund.

## AUTHOR CONTRIBUTIONS

Conceptualization and Writing: RML, MIV, WL, and SSM; Data Curation, Formal Analysis, Investigation, Methodology, Software, and Visualization: RML; Funding Acquisition and Supervision: MIV, WL, and SSM.

## COMPETING INTERESTS

The PTEN^-/-^ cells are licensed by Horizon Discovery Ltd. (Cambridge, UK). Dr. Vitolo receives compensation from the sale of these cells. The remaining authors declare no conflict of interest.

## REFERENCES

1. Cheung, K. J. et al. Polyclonal breast cancer metastases arise from collective dissemination of keratin 14-expressing tumor cell clusters. Proc. Natl. Acad. Sci. 113, E854–E863 (2016).

2. Al Habyan, S., Kalos, C., Szymborski, J. & McCaffrey, L. Multicellular detachment generates metastatic spheroids during intra-abdominal dissemination in epithelial ovarian cancer. Oncogene (2018) doi: 10.1038/s41388-018-0317-x.

3. Alexander, S., Koehl, G. E., Hirschberg, M., Geissler, E. K. & Friedl, P. Dynamic imaging of cancer growth and invasion: a modified skin-fold chamber model. Histochem. Cell Biol. 130, 1147–1154 (2008).

4. Zajac, O. et al. Tumour spheres with inverted polarity drive the formation of peritoneal metastases in patients with hypermethylated colorectal carcinomas. Nat. Cell Biol. (2018) doi: 10.1038/s41556-017-0027-6.

5. Aceto, N. et al. Circulating Tumor Cell Clusters Are Oligoclonal Precursors of Breast Cancer Metastasis. Cell 158, 1110–1122 (2014).

6. Allen, T. A. et al. Circulating tumor cells exit circulation while maintaining multicellularity, augmenting metastatic potential. J. Cell Sci. 132, jcs231563 (2019).

7. Au, S. H. et al. Clusters of circulating tumor cells traverse capillary-sized vessels. Proc. Natl. Acad. Sci. 113, 4947–4952 (2016).

8. Friedl, P., Locker, J., Sahai, E. & Segall, J. E. Classifying collective cancer cell invasion. Nat. Cell Biol. 14, 777–783 (2012).

9. Yang, J. et al. Guidelines and definitions for research on epithelial–mesenchymal transition. Nat. Rev. Mol. Cell Biol. 1–12 (2020) doi: 10.1038/s41580-020-0237-9.

10. Williams, E. D., Gao, D., Redfern, A. & Thompson, E. W. Controversies around epithelial–mesenchymal plasticity in cancer metastasis. Nat. Rev. Cancer 1–17 (2019) doi: 10.1038/s41568-019-0213-x.

11. Kröger, C. et al. Acquisition of a hybrid E/M state is essential for tumorigenicity of basal breast cancer cells. Proc. Natl. Acad. Sci. 116, 7353–7362 (2019).

12. Liu, X. et al. Epithelial-type systemic breast carcinoma cells with a restricted mesenchymal transition are a major source of metastasis. Sci. Adv. 5, eaav4275 (2019).

13. Yu, M. et al. Circulating Breast Tumor Cells Exhibit Dynamic Changes in Epithelial and Mesenchymal Composition. Science 339, 580–584 (2013).

14. Ilina, O. et al. Intravital microscopy of collective invasion plasticity in breast cancer. Dis. Model. Mech. 11, dmm034330 (2018).

15. Mitchel, J. A. et al. The unjamming transition is distinct from the epithelial-to-mesenchymal transition. bioRxiv 665018 (2019) doi: 10.1101/665018.

16. Kim, J. H. et al. Unjamming and collective migration in MCF10A breast cancer cell lines. Biochem. Biophys. Res. Commun. 521, 706–715 (2020).

17. Palamidessi, A. et al. Unjamming overcomes kinetic and proliferation arrest in terminally differentiated cells and promotes collective motility of carcinoma. Nat. Mater. 18, 1252–1263 (2019).

18. Haeger, A., Krause, M., Wolf, K. & Friedl, P. Cell jamming: Collective invasion of mesenchymal tumor cells imposed by tissue confinement. Biochim. Biophys. Acta (2014) doi: 10.1016/j.bbagen.2014.03.020.

19. López-Knowles, E. et al. PI3K pathway activation in breast cancer is associated with the basal-like phenotype and cancer-specific mortality. Int. J. Cancer 126, 1121–1131 (2010).

20. Depowski, P. L., Rosenthal, S. I. & Ross, J. S. Loss of Expression of the PTEN Gene Protein Product Is Associated with Poor Outcome in Breast Cancer. Mod. Pathol. 14, 672–676 (2001).

21. Herbst, R. S. & Schlessinger, J. Small molecule combats cancer-causing KRAS protein at last. Nature 575, 294–295 (2019).

22. Sanchez-Vega, F. et al. Oncogenic Signaling Pathways in The Cancer Genome Atlas. Cell 173, 321-337.e10 (2018).

23. Thompson, K. N. et al. The combinatorial activation of the PI3K and Ras/MAPK pathways is sufficient for aggressive tumor formation, while individual pathway activation supports cell persistence. Oncotarget 6, 35231–46 (2015).

24. Yankaskas, C. L. et al. A microfluidic assay for the quantification of the metastatic propensity of breast cancer specimens. Nat. Biomed. Eng. 1 (2019) doi: 10.1038/s41551-019-0400-9.

25. Lee, R. M., Stuelten, C. H., Parent, C. A. & Losert, W. Collective cell migration over long time scales reveals distinct phenotypes. Converg. Sci. Phys. Oncol. 2, 025001 (2016).

26. Lee, R. M., Kelley, D. H., Nordstrom, K. N., Ouellette, N. T. & Losert, W. Quantifying stretching and rearrangement in epithelial sheet migration. New J. Phys. 15, 025036 (2013).

27. Stuelten, C. H., Lee, R. M., Losert, W. & Parent, C. A. Lysophosphatidic acid regulates the motility of MCF10CA1a breast cancer cell sheets via two opposing signaling pathways. Cell. Signal. 45, 1–11 (2018).

28. Weiger, M. C. et al. Real-Time Motion Analysis Reveals Cell Directionality as an Indicator of Breast Cancer Progression. PLoS One 8, e58859 (2013).

29. C. Kozma, S. et al. The human c-Kirsten ras gene is activated by a novel mutation in codon 13 in the breast carcinoma cell line MDA-MB231. Nucleic Acids Res. 15, 5963–5971 (1987).

30. Eckert, L. B. et al. Involvement of Ras Activation in Human Breast Cancer Cell Signaling, Invasion, and Anoikis. Cancer Res. 64, 4585–4592 (2004).

31. Adeyinka, A. et al. Activated Mitogen-activated Protein Kinase Expression during Human Breast Tumorigenesis and Breast Cancer Progression. Clin. Cancer Res. 8, 1747–1753 (2002).

32. Malaney, S. & Daly, R. J. The Ras Signaling Pathway in Mammary Tumorigenesis and Metastasis. J. Mammary Gland Biol. Neoplasia 6, 101–113 (2001).

33. Cox, T. R. & Erler, J. T. Remodeling and homeostasis of the extracellular matrix: implications for fibrotic diseases and cancer. Dis. Model. Mech. 4, 165–178 (2011).

34. Conklin, M. W. et al. Aligned collagen is a prognostic signature for survival in human breast carcinoma. Am. J. Pathol. 178, 1221–1232 (2011).

35. Padmanaban, V. et al. E-cadherin is required for metastasis in multiple models of breast cancer. Nature 573, 439–444 (2019).

36. Meirson, T., Gil-Henn, H. & Samson, A. O. Invasion and metastasis: the elusive hallmark of cancer. Oncogene 1–3 (2019) doi: 10.1038/s41388-019-1110-1.

37. Hanahan, D. & Weinberg, R. A. Hallmarks of Cancer: The Next Generation. Cell 144, 646–674 (2011).

38. Vitolo, M. I. et al. Deletion of PTEN promotes tumorigenic signaling, resistance to anoikis, and altered response to chemotherapeutic agents in human mammary epithelial cells. Cancer Res. 69, 8275–8283 (2009).

39. Martin, S. S. et al. A cytoskeleton-based functional genetic screen identifies Bcl-xL as an enhancer of metastasis, but not primary tumor growth. Oncogene 23, 4641–4645 (2004).

40. Nair, N. U. et al. Migration rather than proliferation transcriptomic signatures are strongly associated with breast cancer patient survival. Sci. Rep. 9, 10989 (2019).

41. Georgess, D. et al. Twist1-Induced Epithelial Dissemination Requires Prkd1 Signaling. Cancer Res. 80, 204–218 (2020).

42. Lee, R. M. et al. Quantifying topography-guided actin dynamics across scales using optical flow. Mol. Biol. Cell mbc.E19-11-0614 (2020) doi: 10.1091/mbc.E19-11-0614.

43. Chen, S. et al. Actin Cytoskeleton and Focal Adhesions Regulate the Biased Migration of Breast Cancer Cells on Nanoscale Asymmetric Sawteeth. ACS Nano 13, 1454–1468 (2019).

44. Karagiannis, G. S., Condeelis, J. S. & Oktay, M. H. Chemotherapy-induced metastasis in breast cancer. Oncotarget 8, 110733–110734 (2017).

45. Paris, S. dijsktra path finder. MATLAB Central File Exchange https://www.mathworks.com/matlabcentral/fileexchange/17385 (2020).

46. Sveen, J. K. An introduction to MatPIV v. 1.6.1. (2004).

47. Shadden, S. C., Lekien, F. & Marsden, J. E. Definition and properties of Lagrangian coherent structures from finite-time Lyapunov exponents in two-dimensional aperiodic flows. Phys. Nonlinear Phenom. 212, 271–304 (2005).

48. Marel, A.-K. et al. Flow and Diffusion in Channel-Guided Cell Migration. Biophys. J. 107, 1054–1064 (2014).

49. Bazellières, E. et al. Control of cell-cell forces and collective cell dynamics by the intercellular adhesome. Nat. Cell Biol. 17, 409–420 (2014).

