## Supplementary Information for "Distinct Roles of Tumor-Associated Mutations in Collective Cell Migration"

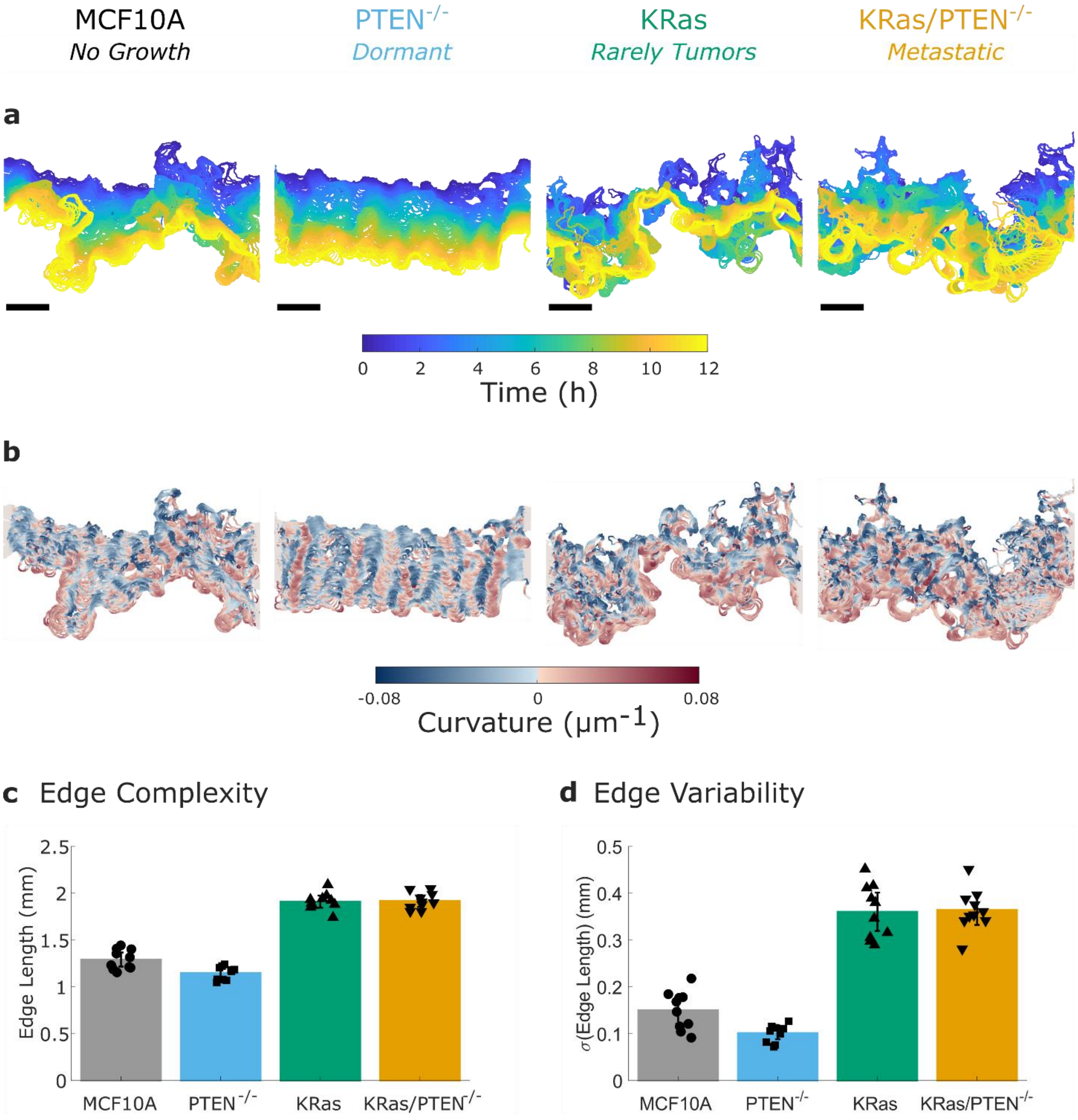


**Supplementary Fig. 1. PTEN$^{-/-}$ and KRas change edge dynamics. a** The dynamics of the leading edge are shown by overlaid edges colored by time. **b** Coloring the leading edge by curvature illustrates the persistence or lack of persistence in local features of the edge shape. **c** Edge length is used to quantify the complexity of the leading edge. **d** The variability in edge length over time is used to quantify the dynamics of the leading edge. N = 10 independent experiments. Error bars indicate 95% confidence interval.

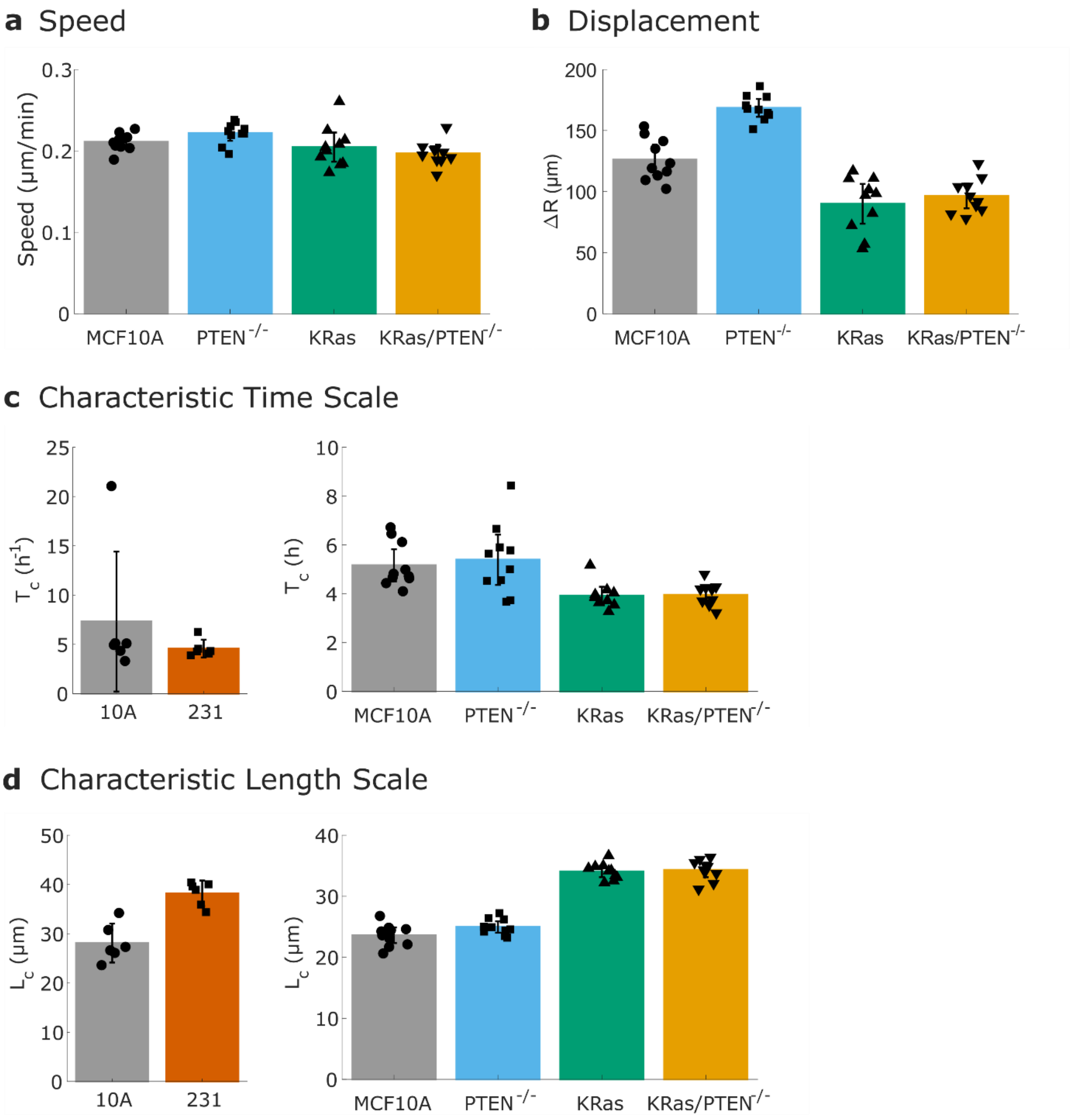


**Supplementary Fig. 2. Additional metrics enhance a multidimensional collective migration phenotype**. **a** Mean speed of the PIV flow field. **b** Displacement of the leading edge. **c** Characteristic time scale of migration calculated using a coarse graining approach. **d** Characteristic length scale of migration calculated using a coarse graining approach. N = 10 (PTEN$^{-/-}$ and KRas) or N = 6 (231) independent experiments. Error bars indicate 95% confidence interval.

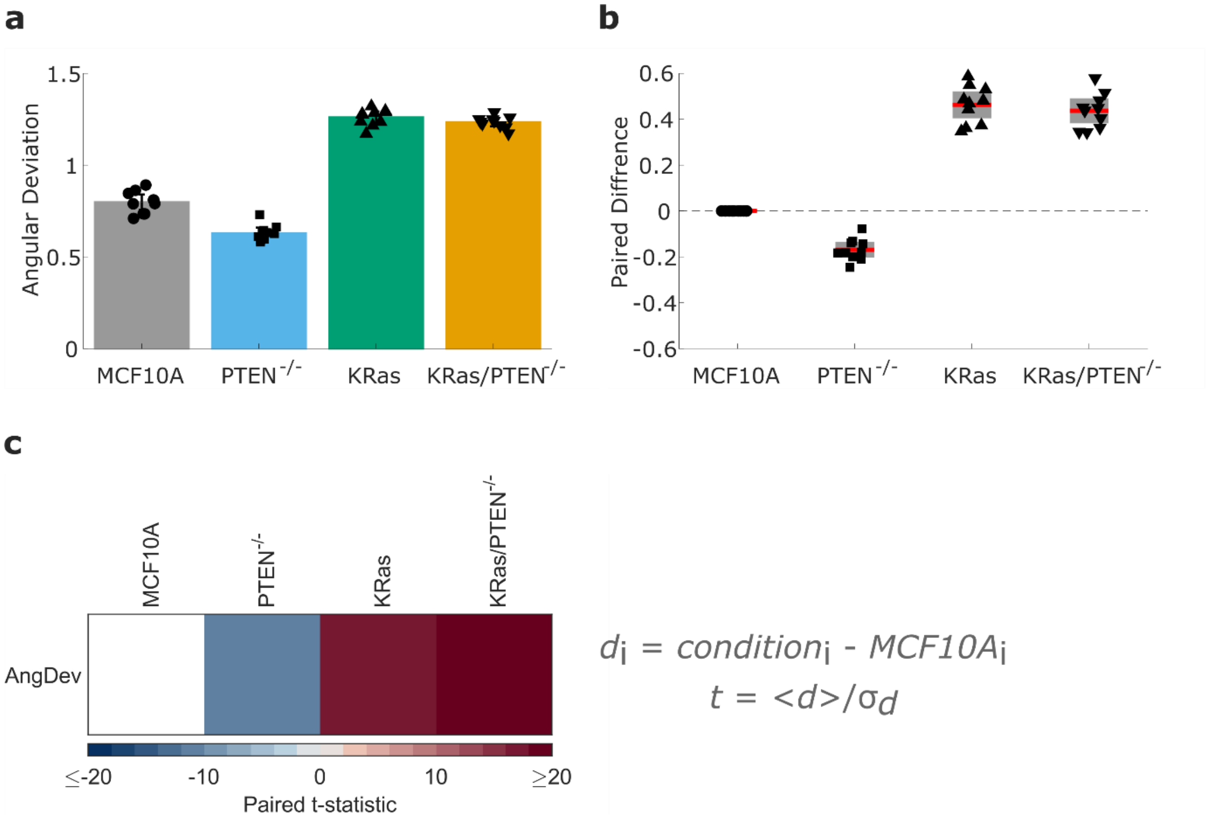


**Supplementary Fig. 3. Paired statistics were used for clustering analysis**. **a** Variability in velocity direction quantified by angular deviation (as shown in **Fig. 3c**). **b** Independent experiments were conducted using paired migration assays, allowing for the calculation of the paired difference for each cell line compared to the same-experiment MCF10A control. **c** These paired differences can be used to calculate to a paired t-statistic. The paired t-statistic is calculated as the mean of the differences divided by the standard deviation of the differences. This allows for the comparison of the strength of changes across metrics which may be measured in different units and on different scales. N = 10 independent experiments. Error bars indicate 95% confidence interval.

### SUPPLEMENTARY MOVIES

**Supplementary Movie 1**. MCF10A (left) and MDA-MB-231 (right) cell sheets migrating over 12 hours. Scale bars are 100 µm and clocks are shown as HH:MM.

**Supplementary Movie 2.** From left to right: MCF10A, $PTEN^{-/-}$, KRas and KRas/$PTEN^{-/-}$ cell sheets migrating over 12 hours. Scale bars are 100 µm and clocks are shown as HH:MM.

### SUPPLEMENTARY DATA

**SupplementaryData.xlsx** contains the source data for all bar graphs, histograms, and correlation curves in Figures 1-3 and the Supplementary Figures, as well as summary tables for Figure 4.
